# Fingering instability accelerates population growth of an expanding cell collective

**DOI:** 10.1101/2023.05.28.542614

**Authors:** Yiyang Ye, Jie Lin

## Abstract

During the expansion of a cell collective, such as the development of microbial colonies and tumor progression, the local cell growth increases the local pressure, which in turn suppresses cell growth. How this pressure-growth coupling affects the expansion of a cell collective remains unclear. Here, we answer this question using a continuum model of cell collective. We find that a fast-growing leading front and a slow-growing interior of the cell collective emerge due to the pressure-dependent growth rate. The leading front can exhibit fingering instability and we confirm the predicted instability criteria numerically with the leading front explicitly simulated. Intriguingly, we find that fingering instability is not only a consequence of local cell growth but also enhances the entire population’s growth rate as positive feedback. Our work unveils the fitness advantage of fingering formation quantitatively and suggests that the ability to form protrusion can be evolutionarily selected.

---

Expanding cell collectives are ubiquitous, e.g., developing microbial colonies, wound healing, and tumor invasions [1–5]. During the expansion, the local pressure and single-cell growth rate can be spatially heterogeneous across the cell collective. For example, an expanding cell collective often displays a leading front underneath which cells actively grow and divide, followed by a crowded interior where cells grow slowly [4–9]. Experiments demonstrated that the low growth rates of interior cells could result from high pressure because a high confining pressure can slow or even completely halt the cell cycle progression [10–12]. Quantitatively, the single-cell growth rate decreases monotonically with its confining pressure due to macromolecular crowding [13, 14].

On the other hand, the pressure itself is generated by active cell growth as a high cell density due to cell division inevitably increases the local pressure. Therefore, the temporal and spatial patterns of pressure and local growth rate are intimately coupled. However, how the interconnection between pressure and cell growth affects the expansion of a cell collective is not understood. In the meantime, the collective’s leading front often exhibits protrusions due to fingering instability [1, 15–18]. While the conditions of fingering instability have been widely studied in different contexts [19–28], it is unclear whether the protrusion can speed up the invasion of a cell collective to its environment relative to a collective without protrusions. If the answer is yes, it is also unclear how to quantify the growth advantage of protrusion formation.

This work combines theoretical analysis and computer simulations to study a continuum model of cell collective in two dimensions. We show that a fast-growing front and a slow-growing interior of the cell collective emerge as an outcome of the pressure-dependent growth rate. The leading front can also exhibit fingering instability, which we derive theoretically. Interestingly, both the pressure gradient and the velocity gradient at the interface contribute to instability. To corroborate with theories, we simulate a numerical model in which the interface separating the cell collective and the environment is explicitly simulated. Surprisingly, the formation of protrusions due to fingering instability enhances the population growth rate by relaxing the pressure and increasing the local growth rate relative to a collective without protrusions under the same parameters. We show that the population growth rate of the collective increases as the surface tension constant decreases. Remarkably, the instability-driven growth enhancement is a consequence of the negative feedback from pressure to growth rate, as we do not see the instability-driven growth enhancement if the local growth rate is strictly constant. Thus, our work provides a minimal model of cell collective expansion incorporating the coupling between pressure and growth rate and a quantitative measure for the fitness advantage of fingering instability.

## The cell-collective expansion model

We study the invasion of a two-dimensional cell collective into a passive fluid (Figure 1). In this section, we ignore the possibility of fingering instability so that the interface is flat. We will discuss the instability condition and its biological impacts later. The collective expands in the *x* direction, and the total length of the system in the *x* direction is *L*. The boundary condition is periodic in the *y* direction with length *L*_*y*_. The velocity field at *x* = 0 is zero by symmetry (Figure 1). We label the *x* coordinate of the interface separating the cell collective and the passive fluid as *x*_*I*_ . The velocity field satisfies Darcy’s law,

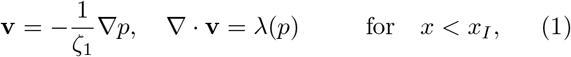

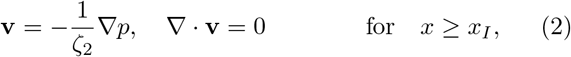

**FIG. 1.**
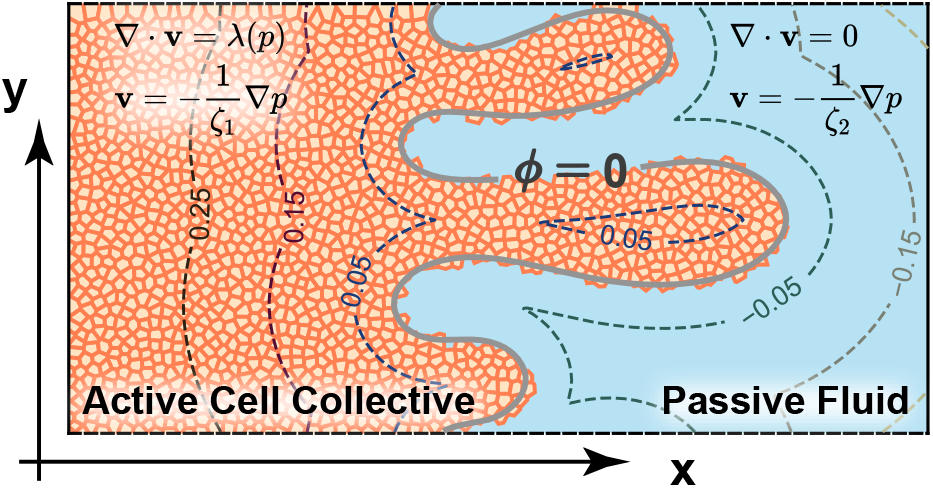
A schematic of a cell collective, in this case, an epithelial tissue, expanding in the *x* direction. In the continuum model, the velocity field is proportional to the pressure gradient. The friction coefficients of the cell collective and the passive fluid can be different. Active growth leads to a finite divergence of the velocity field in the cell collective. In numerical simulations, we use the level-set method to distinguish the cell collective from the passive fluid, and the zero level set (*ϕ* = 0) corresponds to the interface.

where *ζ*_1_ and *ζ*_2_ are the friction coefficients of the cell collective and the passive fluid over the substrate, which can depend on the substrate stiffness [29]. The surrounding fluid is incompressible, while the divergence of the velocity field inside the cell collective is equal to the local growth rate *λ*(*p*), a coarse-grained variable quantifying how fast cells divide locally and a function of local pressure.

Given the pressure-dependence of growth rate and the boundary conditions, which are *∂p*(*x*)*/∂x*|_*x*=0_ = 0 and *p*|_*x*=*L*_ = 0, the pressure field can be uniquely determined assuming a homogeneity along the *y* direction. Here, we set the pressure at the rightmost side of the system to be zero without losing generality since only the pressure gradient matters. To obtain insights from analytical analysis, we set the growth rate to be a linear function of the pressure so that *λ* = *λ*_0_(1 *− p/p*_*c*_) and *λ* = 0 for *p > p*_*c*_. In this linear model, the pressure profile is exactly solvable where

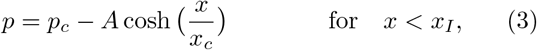

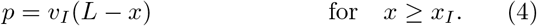

Here, *v*_*I*_ ≡ *v*_*x*_(*x*_*I*_) is the interface velocity, and *A* is a coefficient (see their expressions in the Supplementary Material). They are determined by the continuity <u>conditio</u>n: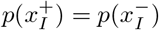, and 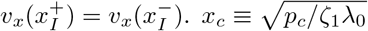 is the characteristic length scale of the pressure fields (Figure 2). Therefore, the local growth rate is

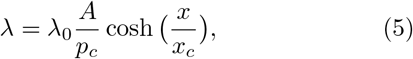

which exhibits a slow-growing region in the collective interior with the characteristic width *x*_*c*_ (Figure 2). In this work, we non-dimensionalize the model so that the length unit is the system size in the *x* direction *L*, the time unit is 1*/λ*_0_, and the pressure unit is *λ*_0_*ζ*_2_*L*^2^. We define the ratio of friction coefficients *α ≡ ζ*_1_*/ζ*_2_. In the following, all variables are dimensionless unless specifically mentioned.

**FIG. 2.**
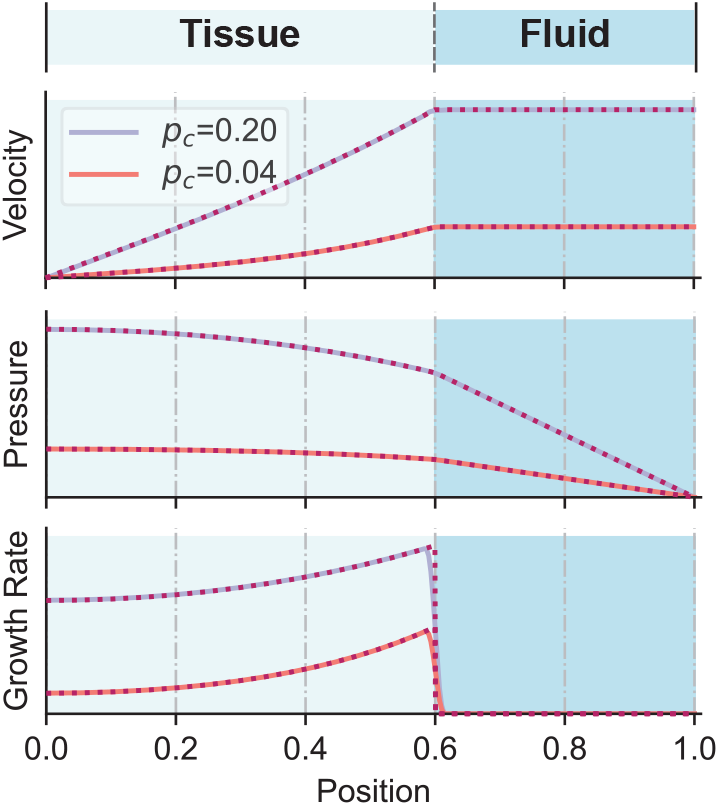
Theoretical predictions in the absence of fingering instability, compared with numerical simulations. The predicted growth rate, pressure, and velocity profiles (dashed lines) agree well with the numerical simulations (solid lines). Cells in the interior of the collective grow slower than cells near the leading front. We show the results of two critical pressures *p*_*c*_.

To study the overall growth of the cell collective, we compute the time derivative of its total volume, *dV*_cell_*/dt*. The population growth is faster when the friction coefficient of the cell collective becomes smaller (Figure S3). Without fingering instability, the surface tension constant of the interface does not enter the spatial patterns of pressure and growth rate and therefore does not affect population growth. Later, we will show that the surface tension constant is crucial in determining the population growth rate due to fingering instability.

### Criteria of fingering instability

In the case of traditional fingering instability, i.e., the Saffman-Taylor instability, the moving interface separating two fluids is pushed by a constant velocity from the infinity [30]. Therefore, only the pressure gradient exhibits a jump across the interface. For an expanding cell collective, the moving interface is pushed by active cell growth. Because of the finite growth rate, the velocity gradient also exhibits a jump across the interface (Figure 2). To test whether both two gradient jumps contribute to interface instability, we introduce a periodic perturbation *ξ*_*k*_(*t*)*e*^i*ky*^ along the interface and derive the time dependence of the perturbation amplitude (Supplementary Material),

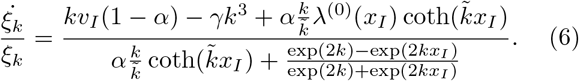

Here, *γ* is the dimensionless surface tension constant with its unit equal to *λ*_0_*ζ*_2_*L*^3^. *λ*^(0)^(*x*_*I*_) is the local growth rate at the leading front in the absence of instability, and 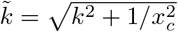. For each interface position *x*_*I*_, the perturbation is unstable within a wave number range (Figure 3A), and we define *k*_*c*_ as the critical wave number below which the interface is unstable against perturbation.

**FIG. 3.**
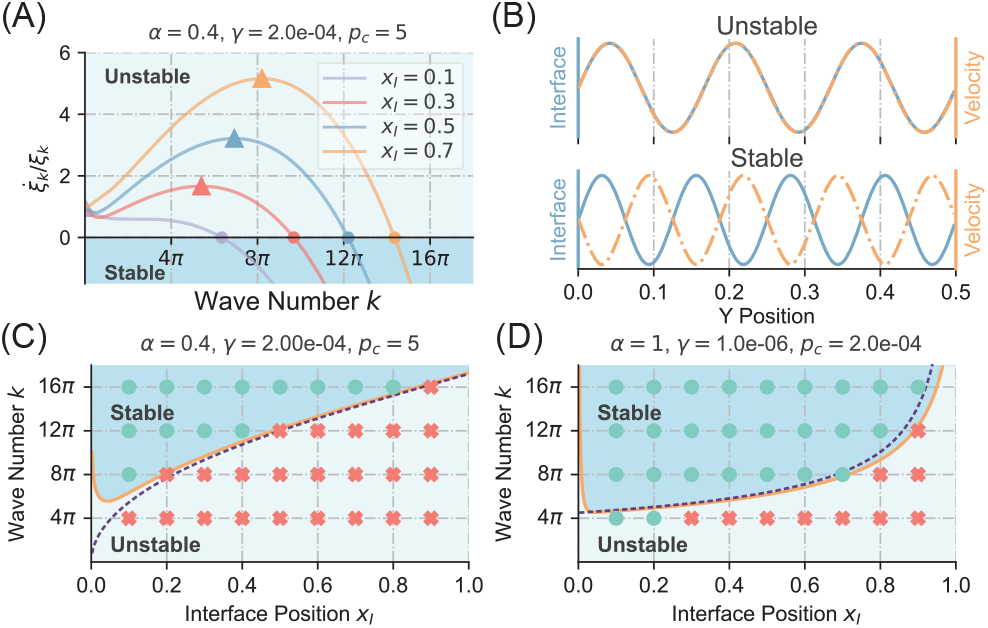
Conditions of fingering instability and numerical verifications. (A) In the linear instability analysis, instability occurs when 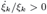. Circles mark the critical wave numbers below which the interface becomes unstable. Triangles mark the wave numbers with the maximum growth rates of the perturbation *ξ*_*k*_. (B) A sinusoidal perturbation is applied to the flat interface in the numerical simulations. A negative correlation between the perturbation and the resulting interface velocity change leads to a stable interface and vice-versa for an unstable interface. The stable example corresponds to *x*_*I*_ = 0.7, *k* = 12*π* in (C), and the unstable example corresponds to *x*_*I*_ = 0.7, *k* = 16*π* in (C). (C and D) Phase diagram for the interface stability. The solid line represents the theoretical result of the linear instability analysis. According to simulations, stable interfaces are marked as green dots, while unstable ones are marked as red crosses. In (C), the interface instability is dominated by the pressure gradient difference across the interface, and the dashed line is the prediction merely considering the effects of the pressure gradient. In (D), the interface instability is purely induced by active cell growth since *α ≡ ζ*_1_*/ζ*_2_ = 1. The dashed line represents the asymptotic result in the limit *x*_*I*_ *≫* 1*/k*_*c*_ *≫ x*_*c*_.

When the friction coefficient of cell collective is smaller than the passive fluid, the pressure gradients are different across the interface because of the different friction coefficients, corresponding to the first term in the numerator of Eq. (6). The different velocity gradients across the interface are due to active cell growth, corresponding to the third term in the numerator of Eq. (6). Because 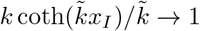 as *k → ∞*, one can always find a *k* so that 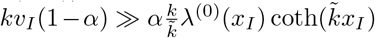 according to Eq. (6). Thus, for a small surface tension constant *γ* 1, the instability is mainly driven by the different pressure gradients. Physically, this is because the small Laplace pressure cannot balance the large pressure drop generated by the distortion of the interface. We confirm this prediction from the phase diagram of interface stability (Figure 3C) where the solid line is the full prediction, and the dashed line is the prediction <u>solely consid</u>ering the pressure gradient, which is 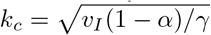.

Interestingly, in the case of active cell collective, instability can be driven by activity alone [25]: even if *α* = 1, the perturbation can still be unstable according to Eq. (6). In the following, we discuss the asymptotic case in which *x*_*I*_ *≫* 1*/k*_*c*_ *≫ x*_*c*_. In this regime, the small *p*_*c*_ imposes a strong constraint on the growth rate, and most cell growth occurs underneath the interface. The growth rate at the interface *λ*_*I*_ = *x*_*c*_*/*(1 *− x*_*I*_), 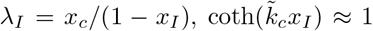, and 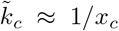 (Supplementary Material). Therefore, the critical wave number 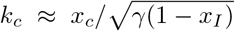, which matches the full prediction pretty well (Figure 3D). Discussions of another asymptotic regime is included in the Supplementary Material, where *p*_*c*_ imposes a negligible effect on the growth rate.

### Simulations of the cell collective expansion

We simulate the expansion of a cell collective by explicitly simulating the interface using the level set method [31–33].

The key idea is to define a level set function *ϕ* as the signed distance to the interface satisfying the constraint |*∇ϕ*| = 1. In our notation, positive *ϕ* represents the cell collective phase, while negative *ϕ* represents the passive fluid phase. The interface is the contour line with *ϕ* = 0 (Figure 1). The constraint |*∇ϕ*| = 1 ensures that if one starts from any point on a contour line of *ϕ* = *ϕ*_0_ and moves along the local normal direction to reach another isoline of *ϕ* = *ϕ*_0_ + *δϕ*, the distance is always equal to *δϕ*. Eqs. (1, 2) are unified using the following equation,

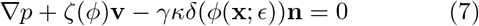

Here, **n** = *−∇ϕ* is the unit vector normal to the interface, and *κ* = *∇*. **n** is the local curvature of the interface. The last term on the left side of Eq. (7) represents the Laplace pressure. *ζ*(*ϕ*) = *ζ*_2_ + (*ζ*_1_ − *ζ*_2_)*H*(*ϕ*; *ϵ*) is the friction coefficient that changes sharply and smoothly across the cell-fluid interface through a smoothed Heaviside function with a thickness 2*ϵ* (Supplementary Material). *δ*(*ϕ*(**x**; *ϵ*)) is a semi-delta function defined as *δ*(*ϕ*; *ϵ*) *≡* d*H*(*ϕ*; *ϵ*)*/* d*ϕ*. As the interface moves, *ϕ* is convected with the velocity field to capture the movement of the interface, 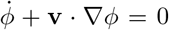. After each time step, we implement a reinitialization process to satisfy the constraint |*∇ϕ* | = 1 using an upwind scheme [31] (Supplementary Material). This constraint ensures that the interface maintains a constant thickness 2*ϵ* during the movement. To verify the validity of our simulations, we compare the simulated pressure and velocity profiles with the theoretical predictions and find that they agree very well (Figure 2).

### Fingering instability accelerates population growth

We next study the biological impacts of fingering instability during the expansion of cell collective. Using numerical simulations, we verify the instability criteria by adding a prescribed perturbation with wave vector *k* along the *y* direction. The perturbation grows if the original perturbation and its following change have a zero phase shift and shrinks if the phase shift is *π* (Figure 3B). We find that Eq. (6) accurately predicts whether the interface is stable or not (Figure 3C, D).

We compare two expanding cell collectives with identical *α* and *p*_*c*_ but different surface tension constants *γ* to unveil the evolutionary advantage of fingering instability. In the cell collective with a small *γ*, we apply a small perturbation to the initial flat interface to trigger fingering instability (Eq. (S28) in the Supplementary Material). In the other collective with a large *γ*, we do not add noise to ensure the interface is flat throughout the simulation (Figure 4A, B). Interestingly, given the same simulation time, the cell collective exhibiting instability grows faster. In particular, the local growth rate and velocity magnitude are much higher in the fingers (Figure 4A, C). In cases of small surface tension constants, we can even observe bubbles made of leader cells disconnected from the bulk (Figure S4). This result suggests that the protrusions can relax the pressure and increase the local growth rate. Indeed, the pressure field in the collective with instability is significantly lower than the stable one (Figure 4B, D).

**FIG. 4.**
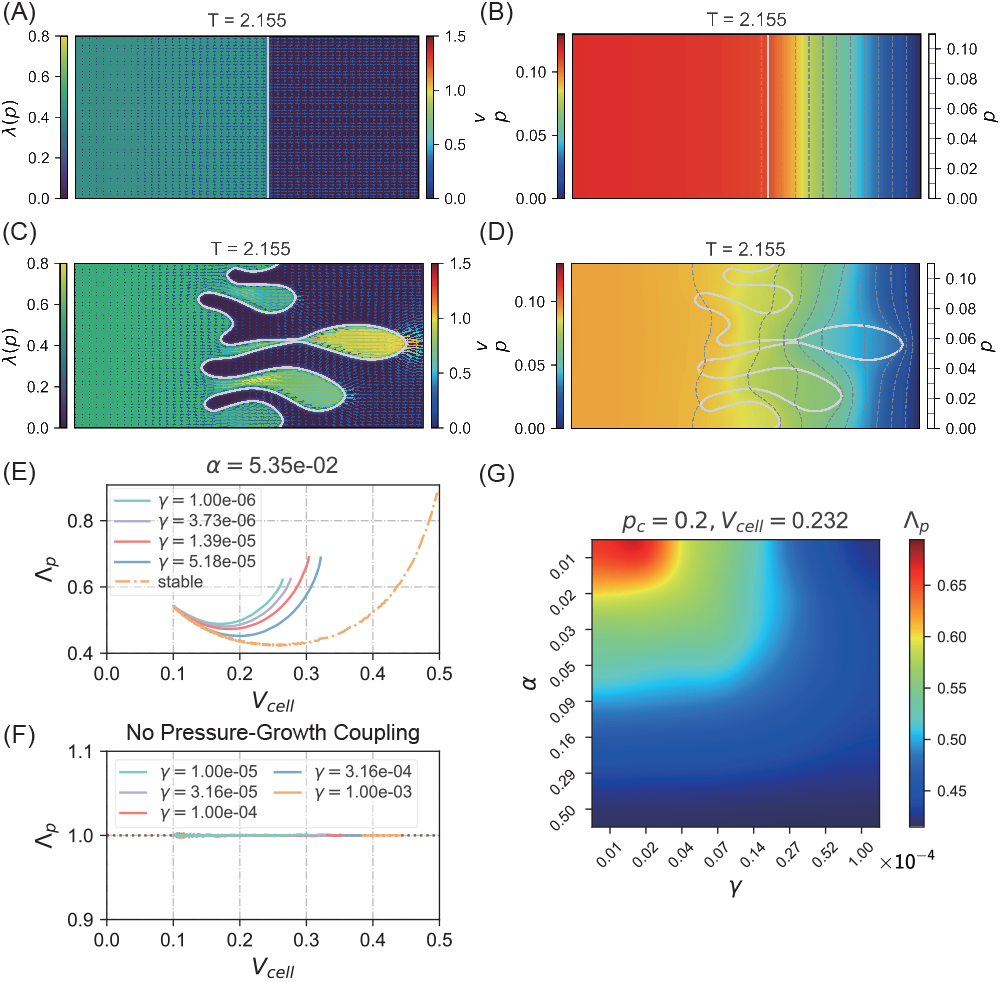
Fingering instability accelerates population growth. (A) The local growth rate pattern of a cell collective at a given time without instability, and the color bar of the growth rate is shown on the left. The arrows represent the velocity field, and the color bar of the velocity magnitude is shown on the right. Here, *α* = 9.35 *×* 10^*−*2^, *γ* = 1 *×* 10^*−*3^. The interface is highlighted as the gray curve. (B) The pressure pattern of the same simulation as (A). The contour lines of the pressure field are shown with the pressure values on the right. The pressure value decreases from left to right. (C) In contrast to (A), we use a small *γ* = 3.73 *×* 10^*−*6^ and apply a perturbation to the initial flat interface so that the cell collective exhibits fingering instability. In this case, the collective grows faster; in particular, the leader cells in the fingers have much higher growth rates and much larger velocity magnitudes. (D) The pressure pattern of the same simulation as (C), which is significantly lower than the case without instability, as shown in (A). (E) The population growth rate of the cell collective Λ_*p*_ as a function of the current volume given different surface tension constants. Each curve is averaged by three independent simulations. (F) The population growth rate of a cell collective without pressure-growth coupling does not depend on *γ* and is always equal to the local cell growth rate Λ_*p*_ = 1. (G) Heatmap of the population growth rate Λ_*p*_ at a given total cell volume *V*_cell_ = 0.232 with respect to *α* and *λ*.

To quantify the advantage of instability, we simulate cell collectives with different surface tension constants *γ* and compute the population growth rate, defined as 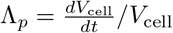. According to Eq. (6), a smaller surface tension constant lets the instability occur earlier. Therefore, we expect cell collectives with smaller *γ* to grow faster. Indeed, the population growth rate increases as the surface tension constant decreases given the same total volume (Figure 4E, G). We also plot the curve of Λ_*p*_ v.s *V*_cell_ without fingering instability as a control to compare (the dotdash line in Figure 4E). We note that in the presence of fingering instability, Λ_*p*_ is a function of both the relative friction coefficient *α* and surface tension constant *γ* (Figure 4G), while in the absence of instability, Λ_*p*_ is only a function of *α* (Figure S5).

We speculate that the acceleration of population growth due to the protrusion formation results from the pressure-growth coupling. To confirm this, we also simulate a modified model in which the growth rate is strictly constant (*λ* = const) and find that, in this case, the population growth rate is always equal to *λ* regardless of the surface tension constant and the morphologies of the cell collective (Figure 4F and Figure S6). This observation shows that the pressure feedback to growth rate is critical to unveil the biological significance of fingering instability.

## Conclusions

In this work, we introduce a minimal model of cell collective in which active cell growth is explicitly included. We incorporate negative feedback from pressure to cell growth, assuming a linear pressure dependence of the local growth rate to make the model analytically solvable. We derive the conditions of fingering instability triggered by the pressure gradient difference and the velocity gradient difference across the interface. To corroborate our theories, we simulate a numerical model in which the interface separating the cell collective and passive fluid is explicitly simulated. All of our theoretical predictions are nicely confirmed. We demonstrate that fingering instability can relieve the pressure confinement on cell growth and let the collective grow faster, particularly for the leader cells in the fingers. Our work provides the first quantitative measure of the fitness advantage of an unstable leading front, which is not only a mechanical consequence of local cell growth but also provides positive feedback to population growth. Therefore, selection pressures on the formation of protrusions may affect the physical properties of cells. For example, the cell surface can be more hydrophilic in an aqueous environment to reduce the surface tension constant.

We thank Lingyu Meng and Sheng Mao for helpful discussions about this work. The research was funded by the National Key R&D Program of China (2021YFF1200500) and supported by Peking-Tsinghua Center for Life Sciences grants.

