## Supplementary Material for "Fingering instability accelerates population growth of an expanding cell collective"

Yiyang Ye[1] and Jie Lin[1,2]
[1] *Center for Quantitative Biology, Peking University, Beijing, China*
[2] *Peking-Tsinghua Center for Life Sciences, Peking University, Beijing, China*
(Dated: May 28, 2023)

### Analytical solutions for a stable cell collective without instability

In this section, we derive the analytical solutions of our model in the absence of fingering instability and use the same normalization convention as in the main text. The length unit is the system length $L$ in the $x$ direction, the pressure unit is $\lambda_0\zeta_2 L^2$, and the time unit is $1/\lambda_0$. Therefore, Eqs. (1, 2) in the main text can be simplified as

$$\begin{aligned} \frac{\partial^2 p_1}{\partial x^2} &= -\alpha\left(1-\frac{p_1}{p_c}\right), \\ \frac{\partial^2 p_2}{\partial x^2} &= 0. \end{aligned} \tag{S1}$$

Here the lower index $i=1$ represents the cell collective, and $i=2$ represents the passive fluid. We label the position of the flat interface as $x_I$, and the pressure field satisfies the following boundary conditions for the cell collective and passive fluid:

$$\left.\frac{\partial p_1}{\partial x}\right|_{x=0} = 0, \qquad \left.\frac{\partial p_1}{\partial x}\right|_{x=x_I^-} = -\alpha v_I, \tag{S2a}$$

$$p_2|_{x=1} = 0, \qquad \left.\frac{\partial p_2}{\partial x}\right|_{x=x_I^+} = -v_I, \tag{S2b}$$

$$p_1|_{x=x_I^-} = p_2|_{x=x_I^+}. \tag{S2c}$$

Here, we consider a left-right symmetrical cell collective expansion so that the velocity at $x=0$ is zero by construction. In addition, the pressure is continuous across a flat interface Eq. (S2c). It is straightforward to find the general solution to Eq. (S1):

$$\begin{aligned} p(x) &= p_c - A\cosh\left(\frac{x}{\sqrt{p_c/\alpha}}\right) \qquad \text{for} \quad x < x_I, \\ p(x) &= v_I(1-x) \qquad \text{for} \quad x \geq x_I. \end{aligned} \tag{S3}$$

Plugging Eq. (S3) into the boundary conditions Eq. (S2), we obtain the coefficients $A$ and $v_I$,

$$\begin{aligned} A &= \frac{\alpha p_c}{\frac{(1-x_I)}{\sqrt{p_c/\alpha}}\sinh\left(\frac{x_I}{\sqrt{p_c/\alpha}}\right) + \alpha\cosh\left(\frac{x_I}{\sqrt{p_c/\alpha}}\right)}, \\ v_I &= \frac{A}{\alpha\sqrt{p_c/\alpha}}\sinh\left(\frac{x_I}{\sqrt{p_c/\alpha}}\right). \end{aligned} \tag{S4}$$

The local growth rate $\lambda(x)$ changes according to the pressure field with a characteristic length scale $x_c = \sqrt{p_c/\alpha}$,

$$\lambda(x) = 1 - \frac{p(x)}{p_c} = \frac{A}{p_c}\cosh\left(\frac{x}{\sqrt{x_c}}\right) \quad \text{for} \quad x < x_I. \tag{S5}$$

### LINEAR INSTABILITY ANALYSIS

In this section, we derive the condition of fingering instability and use the same unit convention as the previous section. Therefore, the unit of the surface tension constant $\gamma$ is $\lambda_0\zeta_2 L^3$. We introduce a perturbation of wavenumber $k$ to the interface,

$$\Gamma_I = x_I + \epsilon \xi_k \exp(ik y), \tag{S6}$$

where $x_I$ is the location of the flat interface and $\xi_k$ is the time-dependent amplitude of the perturbation. The general solution to Eq. (S1) becomes

$$p_i = p_i^{(0)} + \epsilon p_i^{(1)} + O(\epsilon^2) \quad \text{where} \quad i = 1, 2. \tag{S7}$$

Here, $\epsilon$ is a small parameter. The curvature of the interface is $-\frac{\partial^2 \Gamma_I}{\partial y^2}$ so that a positive curvature stands for a bumping pointing from the cell collective into the passive fluid. The pressure field should satisfy the following boundary conditions:

$$\left.\frac{\partial p_1}{\partial x}\right|_{\Gamma_I(y)} = -\alpha \dot{\Gamma}_I(y), \qquad \left.\frac{\partial p_1}{\partial x}\right|_{x=0} = 0, \tag{S8a}$$

$$\left.\frac{\partial p_2}{\partial x}\right|_{\Gamma_I(y)} = -\dot{\Gamma}_I(y), \qquad p_2|_{x=1} = 0, \tag{S8b}$$

$$p_1 - p_2|_{\Gamma_I} = \epsilon\gamma\xi_k k^2 \exp(iky). \tag{S8c}$$

When $\epsilon = 0$, the solution to this system is reduced to $p_i^{(0)}$, which is Eq. (S3). Keeping the first order terms of $O(\epsilon)$, the equations and boundary conditions of $p_i^{(1)}$ become

$$\nabla^2 p_1^{(1)} = \frac{\alpha}{p_c} p_1^{(1)}, \qquad \nabla^2 p_2^{(1)} = 0, \tag{S9a}$$

$$\left.\frac{\partial p_1^{(1)}}{\partial x}\right|_{x_I} = \alpha\left[\xi_k \lambda^{(0)}(x_I) - \dot{\xi}_k\right]\exp(iky), \qquad \left.\frac{\partial p_2^{(1)}}{\partial x}\right|_{x_I} = -\dot{\xi}_k \exp(iky), \tag{S9b}$$

$$\left.\frac{\partial p_1}{\partial x}\right|_{x=0} = 0, \qquad \left.p_2^{(1)}\right|_{x=1} = 0, \tag{S9c}$$

$$\left.p_1^{(1)} - p_2^{(1)}\right|_{x_I} = \left[\gamma k^2 - v_I(1-\alpha)\right]\xi_k \exp(iky). \tag{S9d}$$

Here, $\lambda^{(0)}(x_I) = 1 - p_1^{(0)}(x_I)/p_c$ stands for the growth rate at the interface. We search for the solutions to Eq. (S9a) of the form $p_i^{(1)} = X_i(x)\exp(iky)$ and find that $X_i$ has the following general expression,

$$X_1 = A_1 \exp(-\tilde{k}x) + B_1 \exp(\tilde{k}x),$$
$$X_2 = A_2 \exp(-kx) + B_2 \exp(kx),$$

where $\tilde{k} = \sqrt{k^2 + 1/x_c^2}$. The coefficients $A_1$, $A_2$, $B_1$, $B_2$ can be determined from the boundary conditions in Eqs. (S9b) and (S9c),

$$A_1 = B_1 = \frac{\alpha(\xi_k \lambda^{(0)}(x_I) - \dot{\xi}_k)}{2\tilde{k}\sinh(\tilde{k}x_I)}$$

$$A_2 = \frac{\dot{\xi}_k}{k}\frac{\exp(2k + kx_I)}{\exp(2k) + \exp(2kx_I)}, \quad B_2 = -\frac{\dot{\xi}_k}{k}\frac{\exp(kx_I)}{\exp(2k) + \exp(2kx_I)}.$$

Finally, using the pressure discontinuity condition across the interface Eq. (S9d), we obtain the time dependence of the perturbation amplitude

$$\frac{\dot{\xi}_k}{\xi_k} = \frac{k v_I(1-\alpha) - \gamma k^3 + \alpha\frac{k}{\tilde{k}}\lambda^{(0)}(x_I)\coth(\tilde{k}x_I)}{\alpha\frac{k}{\tilde{k}}\coth(\tilde{k}x_I) + \frac{\exp(2k)-\exp(2kx_I)}{\exp(2k)+\exp(2kx_I)}}, \tag{S10}$$

which is Eq. (6) in the main text.

Since the denominator on the right-hand side of Eq. (S10) is always positive, the instability can be triggered by the difference in the friction coefficients, that is, the $kv_I(1-\alpha)$ term in the numerator. The instability can also be triggered by active growth, the $\lambda^{(0)}(x_I) > 0$ term in the numerator. We define the critical wavenumber $k_c$ as the root of $\dot{\xi}_k = 0$. As we discuss in the main text, when $\alpha < 1$, the instability is caused by the difference in the friction coefficient as long as the surface tension $\gamma$ is small enough. In this case, $k_c = \sqrt{\frac{v_I(1-\alpha)}{\gamma}}$. Unlike the traditional Saffman-Taylor instability, even when $\alpha = 1$, the system can still produce instability through active growth. The critical wavenumber $k_c$ is determined by the following equation

$$\gamma k_c^2 \tilde{k}_c = \lambda^{(0)}(x_I)\coth(\tilde{k}_c x_I). \tag{S11}$$

We find that the root of Eq. (S11) is determined by the following three characteristic lengths, $x_I$, $1/k_c$, and $x_c = \sqrt{p_c}$.

In the following, we discuss two asymptotic cases in which $k_c$ has a simple form. The first situation is $x_c \gg x_I \gg 1/k_c$. $x_c \gg x_I$ implies that pressure feedback has a negligible effect on the growth rate. Therefore, the growth rate of the cell collective at any position is close to 1, which means $\lambda^{(0)}(x_I) \approx 1$. In addition, $\coth(\tilde{k}_c x_I) \approx 1$ holds. Therefore, Eq. (S11) can be reduced to

$$\tilde{k}_c \approx k_c \approx \frac{1}{\sqrt[3]{\gamma}}. \tag{S12}$$

We verify the phase diagram in this asymptotic case by numerical simulations (Fig. S1).

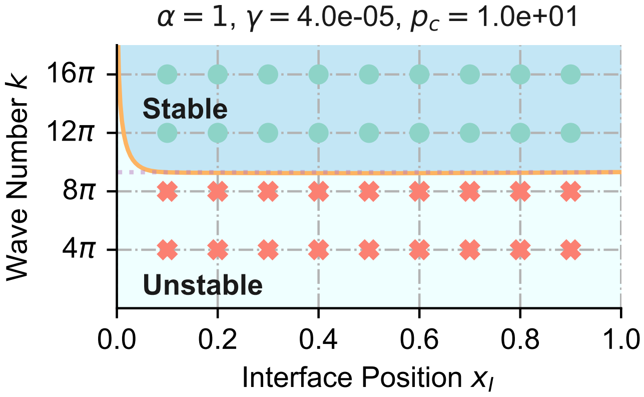


FIG. S1. Phase diagram of interface stability. The solid line represents the theoretical results calculated from Eq. (S11). The dotted line represents the asymptotic result in Eq. (S12). According to the numerical simulations, stable interfaces are marked as green dots, while unstable ones are marked as red crosses.

The second situation is $x_I \gg 1/k_c \gg x_c$. Contrary to the previous situation, $x_I \gg x_c$ means that $p_c$ imposes a strong constraint on the growth rate. In this case, active cell growth is localized underneath the interface. It can be seen from Eq. (S3) cells near the interface grow with the growth rate,

$$\lambda^{(0)}(x_I) = 1 - \frac{p(x_I)}{p_c} = \frac{1}{1 + \frac{1-x_I}{x_c}\tanh\left(\frac{x_I}{x_c}\right)} \approx \frac{x_c}{1-x_I}. \tag{S13}$$

Combined with $\coth(\tilde{k}_c x_I) \approx 1$, and $\tilde{k}_c \approx 1/x_c$, Eq. (S11) can be reduced to

$$k_c \approx \frac{x_c}{\sqrt{\gamma(1-x_I)}}, \tag{S14}$$

which has been verified in Figure 3D in the main text.

### DETAILS OF NUMERICAL SIMULATION

Our numerical simulations use the standard level set approach to represent the moving interface between the cell collective and the passive fluid under a fixed mesh grid [1–3]. We introduce an auxiliary field $\phi$ defined as the signed distance to the interface as the level set function. Physical properties that distinguish the two different phases are functions of $\phi$,

$$\zeta(\phi) = \zeta_2 + (\zeta_1 - \zeta_2)H(\phi;\epsilon)$$
$$\lambda(\phi,p) = \lambda_0(\phi)(1 - p/p_c) = \lambda_0(1 - p/p_c)H(\phi;\epsilon).$$

Here, $H(\phi;\epsilon)$ is a smoothed Heaviside step function of $\phi$ with a transition zone of width $2\epsilon$,

$$H(\phi;\epsilon) = \begin{cases} 0, & \phi < -\epsilon \\ \frac{1}{2}[1 + \frac{\phi}{\epsilon} + \frac{1}{\pi}\sin(\pi\frac{\phi}{\epsilon})], & -\epsilon \leq \phi \leq \epsilon \\ 1, & \phi > \epsilon \end{cases} \tag{S15}$$

$\epsilon$ is five times the simulation grid size in our simulations. With the help of $\phi$, we can write the velocity divergence as

$$\nabla \cdot \mathbf{v} = \lambda(\phi,p), \tag{S16}$$

which simultaneously describes the growth of the cell collective and the incompressibility of the passive fluid. The unit vector normal to the interface is defined as $\mathbf{n} = -\nabla\phi/|\nabla\phi|$, which points from the cell collective (where $\phi > 0$) to the passive fluid (where $\phi < 0$). Therefore, the sign of the curvature $\kappa = \nabla \cdot \mathbf{n}$ follows the same convention as the previous section. In an ideal system with a sharp interface, the pressure has a discrete jump across the interface,

$$p_1 = p_2 + \kappa\gamma. \tag{S17}$$

To write the discontinuous condition Eq. (S17) into a continuous form with a finite interface thickness, we consider an arbitrary control region $\Omega$ across the interface (Fig. S2). Force balance equations for region $\Omega_1$ lying in the cell

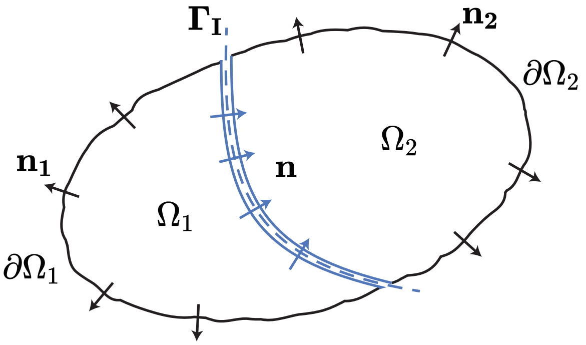


FIG. S2. Analyze the forces acting on a control volume $\Omega$ containing the cell-fluid interface $\Gamma_I$ in order to unify the force equilibrium condition into a single equation as is shown in Eq. (S20).

collective phase and for region $\Omega_2$ lying in the passive fluid phase take the form of

$$\int_{\Omega_1} -\zeta_1 \mathbf{v} dv - \int_{\partial\Omega_1} p\mathbf{n_1} ds - \int_{\Gamma_I} p_1 \mathbf{n} ds = 0, \tag{S18}$$

$$\int_{\Omega_2} -\zeta_2 \mathbf{v} dv - \int_{\partial\Omega_2} p\mathbf{n_2} ds + \int_{\Gamma_I} p_2 \mathbf{n} ds = 0. \tag{S19}$$

Here, $\Gamma_I$ is the interface. Adding Eq. (S18) and Eq. (S19), and using the discontinuous condition Eq. (S17), we get

$$\int_{\Omega} -\zeta(\phi)\mathbf{v} dv - \int_{\Gamma_I} \kappa\gamma\mathbf{n} ds - \sum_i \int_{\partial\Omega_i} p\mathbf{n_i} ds = 0. \tag{S20}$$

We simplify the second term in Eq. (S20) to a volume integral using the smoothed $\delta$-function,

$$\int_{\partial\Gamma_I} \kappa\gamma\mathbf{n} ds = \int_{\Omega} \kappa\gamma\mathbf{n}\, |\nabla\phi|\, \delta(\phi;\epsilon) dv = -\int_{\Omega} \kappa\gamma\nabla\phi\delta(\phi;\epsilon) dv. \tag{S21}$$

Finally, the force balance equation is simplified to

$$\nabla p + \zeta(\phi) v - \underbrace{\kappa\gamma\delta(\phi(\mathbf{x});\epsilon)\nabla\phi}_{f_{\text{surf}}(\phi)} = 0. \tag{S22}$$

The differential equation that the pressure field $p$ satisfies can be obtained by combining the force balance equation Eq. (S22) with the velocity divergence equation Eq. (S16),

$$\nabla^2 p - \frac{\nabla\zeta(\phi)}{\zeta(\phi)} \cdot \nabla p - \frac{\lambda_0(\phi)\zeta(\phi)}{p_c} p + \zeta(\phi)\left[\lambda_0(\phi) - \nabla \cdot \left(\frac{f_{\text{surf}}(\phi)}{\zeta(\phi)}\right)\right] = 0. \tag{S23}$$

In deriving Eq. (S23), we have used

$$\mathbf{v} = \frac{f_{\text{surf}}(\phi) - \nabla p}{\zeta(\phi)}. \tag{S24}$$

Note that Eq. (S23) is a second-order linear partial differential equation of $p$, with coefficients only depending on $\phi$. Therefore, Eq. (S23) can be solved directly after discretization by the central difference method. From the solution of $p$, we immediately obtain $v$ using Eq. (S24). $\phi$ is updated according to the following equation:

$$\frac{\mathcal{D}}{\mathcal{D}t}\phi = \dot{\phi} + \mathbf{v} \cdot \nabla\phi = 0, \tag{S25}$$

where $\mathcal{D}/\mathcal{D}t$ stands for the material derivative. We add a reinitialization process to satisfy the constraint $|\nabla\phi| = 1$ by iterating the following equation for a few "artificial" time step $\mathrm{d}\tau$ after every real-time update of the level set function Eq. (S25).

$$\frac{\partial\phi}{\partial\tau} + S(\phi^{(n,0)};\epsilon)(|\nabla\phi| - 1) = 0, \tag{S26}$$

where $S(\phi^{(n,0)};\epsilon) = 2H(\phi^{(n,0)};\epsilon) - 1$ is a smoothed sign function with a finite thickness $\epsilon$. $S(\phi^{(n,0)};\epsilon)$ is calculated from the initial configuration of $\phi$ after each update Eq. (S25). The first superscript $n$ is served for the number of total iterations over the actual time step $\mathrm{d}t$, while the second superscript indicates the number of iterations over the virtual time step $\mathrm{d}\tau$ at the present moment. Eq. (S26) lets the cell-fluid interface described by $\phi = 0$ remains fixed during the reinitialization process. We rearrange Eq. (S26) as the following advection equation,

$$\frac{\partial\phi}{\partial\tau} + \underbrace{\left(S(\phi^{(n,0)})\frac{\nabla\phi}{|\nabla\phi|}\right)}_{\mathbf{w}} \cdot \nabla\phi = S(\phi^{(n,0)}),$$

where $\mathbf{w}$ is the wind velocity. Since the wind velocity is towards the cell collective and passive fluid from the interface, the reinitialization starts at the interface and moves outwards. Efficient numerical schemes for solving Eq. (S26) were introduced in [4]. The simplest first-order scheme used in our two-dimensional simulation is given by

$$\phi^{(n,m+1)} = \phi^{(n,m)} + \Delta\tau S(\phi^{(n,0)})\left(1 - \sqrt{D_x^2 + D_y^2}\right), \tag{S27}$$

where $D_x$ and $D_y$ are the discrete derivatives of $\phi$ in the $x$ and $y$ direction. They are calculated using a set of data points biased to be more "upwind" of the query point, with respect to the direction of the wind velocity $\mathbf{w}$. Detailed expressions of $D_x$ and $D_y$ are

$$D_x = \begin{cases} D_x^+, & \text{if } w_x^+ < 0 \text{ and } w_x^+ + w_x^- < 0 \\ D_x^-, & \text{if } w_x^- > 0 \text{ and } w_x^+ + w_x^- > 0 \\ 0, & \text{if } w_x^- < 0 \text{ and } w_x^+ > 0 \end{cases}$$

$$D_y = \begin{cases} D_y^+, & \text{if } w_y^+ < 0 \text{ and } w_y^+ + w_y^- < 0 \\ D_y^-, & \text{if } w_y^- > 0 \text{ and } w_y^+ + w_y^- > 0 \\ 0, & \text{if } w_y^- < 0 \text{ and } w_y^+ > 0, \end{cases}$$

where

$$
\begin{aligned}
w_x^+ &= S(\phi^{(n,0)})D_x^+, \qquad & w_x^- &= S(\phi^{(n,0)})D_x^-, \\
w_y^+ &= S(\phi^{(n,0)})D_y^+, \qquad & w_y^- &= S(\phi^{(n,0)})D_y^-, \\
D_x^+ &= \frac{\phi_E - \phi_C}{d}, \qquad & D_x^- &= \frac{\phi_C - \phi_W}{d}, \\
D_y^+ &= \frac{\phi_N - \phi_C}{d}, \qquad & D_y^- &= \frac{\phi_C - \phi_S}{d}.
\end{aligned}
$$

The subscripts E, W, N, and S indicate the grid points in the directions of east, west, north, and south, respectively. We remark that the error of $\phi$ is computed only near the interface since the deviation of $|\nabla\phi|$ from 1 is mostly localized to the interface in our simulation. We summarize our numerical simulation protocol in Algorithm 1.

In the presence of noise, the initial shape of the interface is in the form of superimposing periodic disturbances on a flat interface

$$\Gamma_I = x_I + \sum_{i=1}^{N} A_i \cos\left(\frac{2\pi N y}{L_y} + \psi_i\right). \tag{S28}$$

When examining the stability of an interface under a specific wavenumber $k = 2\pi j/L_y$, we take $A_i = 0.002\delta_{i,j}$ and $\psi_i = 0$ (Figure 3B in the main text). To simulate cell collectives with fingering instability, we impose white noise perturbation on the interface by superposition of 10 sinusoidal functions with the same amplitude $A_i = 0.001$ and random phase $\psi_i \in [0, 2\pi]$ (e.g., Figure 4C in the main text).

**Algorithm1** Level Set Simulation

**Initialization:** set $\phi^{(0,0)}$ as initial condition
**while** $t < t_f$ **do**
  calculate surface tension $f_{\text{surf}}$ from $\phi^{(n,0)}$, see Eq. (S22)
  solve pressure field $p$ from Eq. (S23)
  calculate velocity field $\mathbf{v}$ from Eq. (S24)
  update $\phi^{(n,0)}$ from Eq. (S25)
  calculate the error of $\phi$: $res^{(0)} = \underset{|\phi|<\epsilon}{\text{Mean}}\{|1 - |\nabla\phi^{(n,0)}||\}$
  $i \leftarrow 0$
  **if** $res > trigger$ **then**
    **repeat**
      **Reinitialize iteration:** update $\phi^{(n,i+1)}$ from $\phi^{(n,i)}$ according to Eq. (S27)
      calculate the error of $\phi$: $res^{(i+1)} = \underset{|\phi|<\epsilon}{\text{Mean}}\{|1 - |\nabla\phi^{(n,i+1)}||\}$
      $i \leftarrow i + 1$
    **until** converge: $|res^{(i+1)} - res^{(i)}| < tol$
  **end if**
  $n \leftarrow n + 1$, $t \leftarrow t + \mathrm{d}t$, $\phi^{(n+1,0)} \leftarrow \phi^{(n,i)}$
**end while**


[1] M. Sussman, P. Smereka, and S. Osher, A level set approach for computing solutions to incompressible two-phase flow, Journal of Computational physics **114**, 146 (1994).
[2] M. Sussman and E. Fatemi, An efficient, interface-preserving level set redistancing algorithm and its application to interfacial incompressible fluid flow, SIAM Journal on scientific computing **20**, 1165 (1999).
[3] J. A. Sethian and P. Smereka, Level set methods for fluid interfaces, Annual review of fluid mechanics **35**, 341 (2003).
[4] S. Osher and J. A. Sethian, Fronts propagating with curvature-dependent speed: Algorithms based on hamilton-jacobi formulations, Journal of computational physics **79**, 12 (1988).

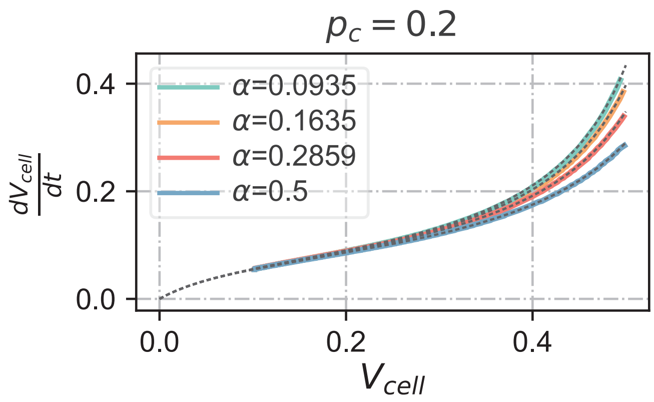


FIG. S3. The growth curve of a growing cell collective with a flat interface. The smaller the friction coefficient, the faster the collective grows.

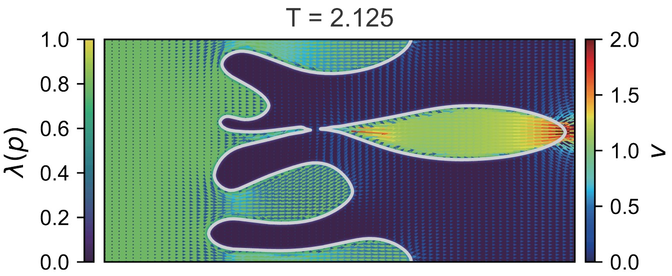


FIG. S4. The level-set simulation method can handle complex interfacial topology during cell collective expansion. A bubble made of leader cells disconnected from the bulk is observed in this simulation. Here $\alpha = 0.05$, $\gamma = 5 \times 10^{-6}$ and $p_c = 0.4$.

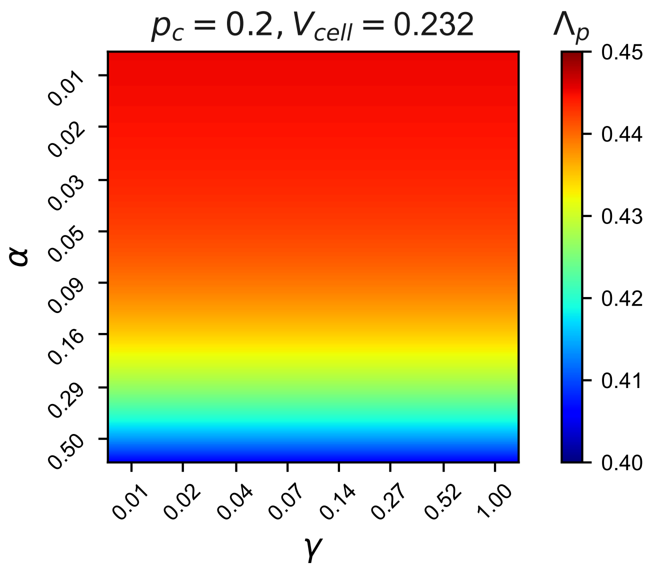


FIG. S5. Heatmap of the population growth rate $\Lambda_p$ with respect to $\alpha$ and $\lambda$ without fingering instability. $\Lambda_p$ is calculated at a given total cell volume $V_{cell} = 0.232$.

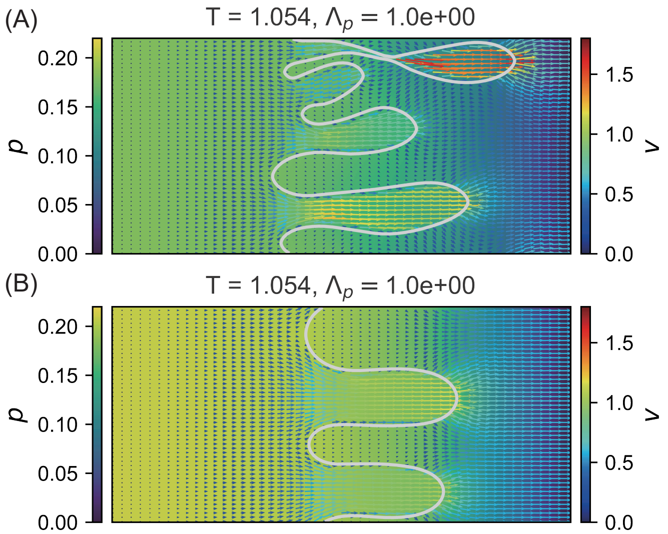


FIG. S6. The pressure patterns and the velocity fields of cell collectives growing without pressure feedback to the local growth rate. A smaller surface tension $\gamma = 10^{-5}$ in (A) produces finer fingers than that in (B) where $\gamma = 10^{-4}$. Though the bulk pressure is reduced due to pronounced instability in (A), the population growth rate is strictly equal to the constant local growth rate in both (A) and (B).
